# Unilateral loss of tyrosine hydroxylase in adult midbrain dopaminergic neurons reveals reduction to neuregulin1 in rat model of acute dopaminergic dysfunction

**DOI:** 10.64898/2026.10.09.757761

**Authors:** Lacey K Greer, Abigail E. Harr, Alexander F Hoffman, Antti J Jӓmsӓ, Mikko Airavaara, Carl R Lupica, Brandon K Harvey

## Abstract

Dopamine, produced primarily in the midbrain, is required for movement and reward-based signaling all throughout life. Loss of dopaminergic function can result in changes to motor systems, mood, motivation, and cognition. Dopaminergic cell death or dopaminergic hypofunction are implicated in a variety of diseases including synucleinopathies, neuropsychiatric disorders, and substance use disorders. Loss of dopaminergic neurons (DANs) in the substantia nigra pars compacta (SNpc) and subsequent dysregulation of dopamine signaling in striatum are implicated to varying degrees in all synucleinopathies. By the time patients present with motor symptoms in Parkinson’s disease, the most common synucleinopathy, >30% of DANs are lost in the SNpc because of ongoing degeneration for years prior to diagnosis. Studying the early consequences of dopamine loss is critical to understanding mechanisms of dopamine depletion disease progression and identifying biomarkers for earlier diagnosis so that effective treatment can ensue. Here, we created a model of dopamine loss in adult rats to study early transcriptional changes following loss of dopamine synthesis. We used CRISPR to knock out tyrosine hydroxylase (TH), the rate-limiting enzyme for dopamine synthesis, in the adult rat midbrain through injection of *Th* guide RNAs into transgenic rats expressing Cas9 nuclease specifically in dopamine transporter positive neurons and performed single nucleus RNA sequencing on midbrain and dorsal striatal tissue. Two weeks after injection, we measured a 90% loss of TH and dopamine in the midbrain, but striatal TH and dopamine were unaffected. Despite preceding striatal dopamine loss, our data identified alterations to genes involved in synaptic signaling, organization, and function in both midbrain and striatal neurons. We found the neurotrophic factor, neuregulin1, to be significantly reduced in dopaminergic neurons, which may account for transcriptional changes in surrounding cells that precede detectable reductions to striatal dopamine. These data establish a new rat model of adult TH loss and identify early transcriptional consequences of dopamine loss in the adult midbrain after normal development, providing potential biomarkers for early-stage dopamine loss in PD synucleinopathies and related disorders of dopamine depletion.

## Introduction

Dopamine (DA) is a neurotransmitter critical for mediating movement and reward signaling^1^. Approximately 90% of dopaminergic neurons (DAN) are located in the midbrain in two key regions: the ventral tegmental area (VTA), involved in reward-based behavior, motivation, and cognition and the substantia nigra pars compacta (SNpc), involved in executing coordinated movement^2,3^. Reduced dopamine signaling in the adult midbrain is a feature in the pathology of neurological conditions such as synucleinopathies, including Parkinson’s disease (PD) and Multiple systems atrophy (MSA), as well as conditions like substance use disorder (SUD)^4,5^. While common pathology of synucleinopathies centers around aggregation of the protein α-synuclein, synucleinopathy-based diseases have at least a “parkinsonian-like” disease subtype where loss of DANs in the SNpc and resulting loss of dopaminergic innervation in the dorsal striatum results in severe motor deficits^6–11^. Substance use disorders such as chronic cocaine use also demonstrate midbrain DAN hypofunction overtime, with less phasic dopamine release in the dorsal and ventral striatum, where midbrain dopaminergic neurons project^12^. Identifying early cellular adaptations to dopamine loss in dopamine depletion diseases may lead to earlier interventional treatments.

Animal models of dopamine depletion have mainly focused on neurotoxin and genetic perturbations for study and developing treatments. Toxins such as 6-hydroxydopamine (6-OHDA) and 1-methyl-4-phenyl-l,2,3,6-tetrahydropyridine (MPTP) have been beneficial for modeling inflammation and DAN death seen in synucleinopathies, but do not accurately represent a gradual dopamine loss as cell death begins within hours of injection^13–16^. Genetic animal models of synucleinopathies express disease relevant mutations in α-synuclein that are causally linked to some form of human synucleinopathies ^17,18^. Although useful, these models do not specifically address the consequences of dopamine depletion that can occur in cases of SUD or idiopathic PD. One popular genetic model of dopaminergic dysfunction is the *Pitx3* ^-/-^(aphakia *or ak)* mouse model in which dopaminergic system fails to develop due to a homozygous mutation in the *Pitx3* homeobox transcription factor, which promotes development, differentiation, and maintenance of midbrain dopaminergic neurons. Studies with aphakia mice demonstrate deficits in motor function and olfaction as well as alterations to striatal medium spiny neuron spiny density, all of which support the model for studying dopamine deficits ^19–21^. However, because the aphakia mouse is a developmental knockout, these animals have never had a fully functional dopaminergic system, in contrast to patients of dopamine depletion diseases, where dopamine signaling is lost in adulthood.

Because PD patients have already lost >30% of their SNpc dopaminergic neurons at the point of diagnosis, it is likely that neurodegeneration has continued for decades prior to onset of motor symptoms ^22,23^. Many cell types in the midbrain and striatum respond to dopamine. Loss of dopamine signaling within dopamine depletion diseases can have effects on a variety of cell types and neurocircuits, beyond striatal medium spiny neurons that are thought to mediate motor and reward signaling^24,25^. Much of the midbrain degeneration research focuses on the later stages of these diseases, when motor symptoms have already occurred, but a better understanding of the early phase of dopamine loss, prior to cell death, could help elucidate early disease symptoms, and identify biomarkers that allow for earlier diagnosis and intervention.

With this in mind, we aimed to identify the early transcriptional changes occurring in the midbrain and striatal cell populations after disrupting the midbrain dopaminergic system of adult rat. We injected Dat-iCre x LSL-Cas9 rats, which express Cas9 nuclease specifically in dopamine transporter (DAT) positive dopaminergic neurons, with guide RNAs (gRNA) specific to tyrosine hydroxylase (TH), the enzyme necessary for dopamine synthesis^26,27^. We performed snRNAseq on midbrain and striatal samples to identify the early transcriptional changes that occur after disrupting dopamine synthesis and found alterations to neuronal signaling pathways and a downregulation of the neurotrophic factor, neuregulin1 just two weeks after injection.

## Results

### Knockout of tyrosine hydroxylase in adult rat midbrain and its impact on dopamine dynamics

To disrupt dopamine synthesis, we injected adeno-associated virus (AAV) expressing gRNAs specific to the *Th* gene into the midbrain of adult Dat-iCre x LSL-Cas9 rats. While Cas9-mediated mutagenesis causes a variety of mutations to create a loss of function, rather than a classical knockout which deletes a consistent gene segment or whole gene, we will be referring to gRNA directed mutagenesis of the *Th* gene as “*Th* knockout”. Animals were contralaterally injected with a control gRNA targeted to the *Rosa26* locus. Both viruses expressed enhanced green fluorescent protein (GFP) as a transduction marker. Two weeks post injection, midbrain TH immunoreactivity was reduced by >90% in the side receiving *Th* gRNAs compared to the control hemisphere (Figure 1A-B). GFP expression is visible, indicating that transduced dopaminergic neurons are still present, despite the reduction in TH levels. To visualize TH loss in the dopaminergic projections, we immunolabeled the dorsal striatum for TH. At two weeks post injection, there was no significant difference between striatal hemispheres (Figure 1C-D). While TH was significantly decreased in the soma, it was unchanged in the processes, likely due to differences in TH turnover rate between the soma and axons ^28^. These data indicate that at this early timepoint of two weeks post injection, we knocked out tyrosine hydroxylase in the midbrain, but striatal TH remained intact. In contrast, TH immunoreactivity was significantly decreased in both midbrain and striatum at 4 weeks post injection (Figure 1 E-H).

**Figure 1.**
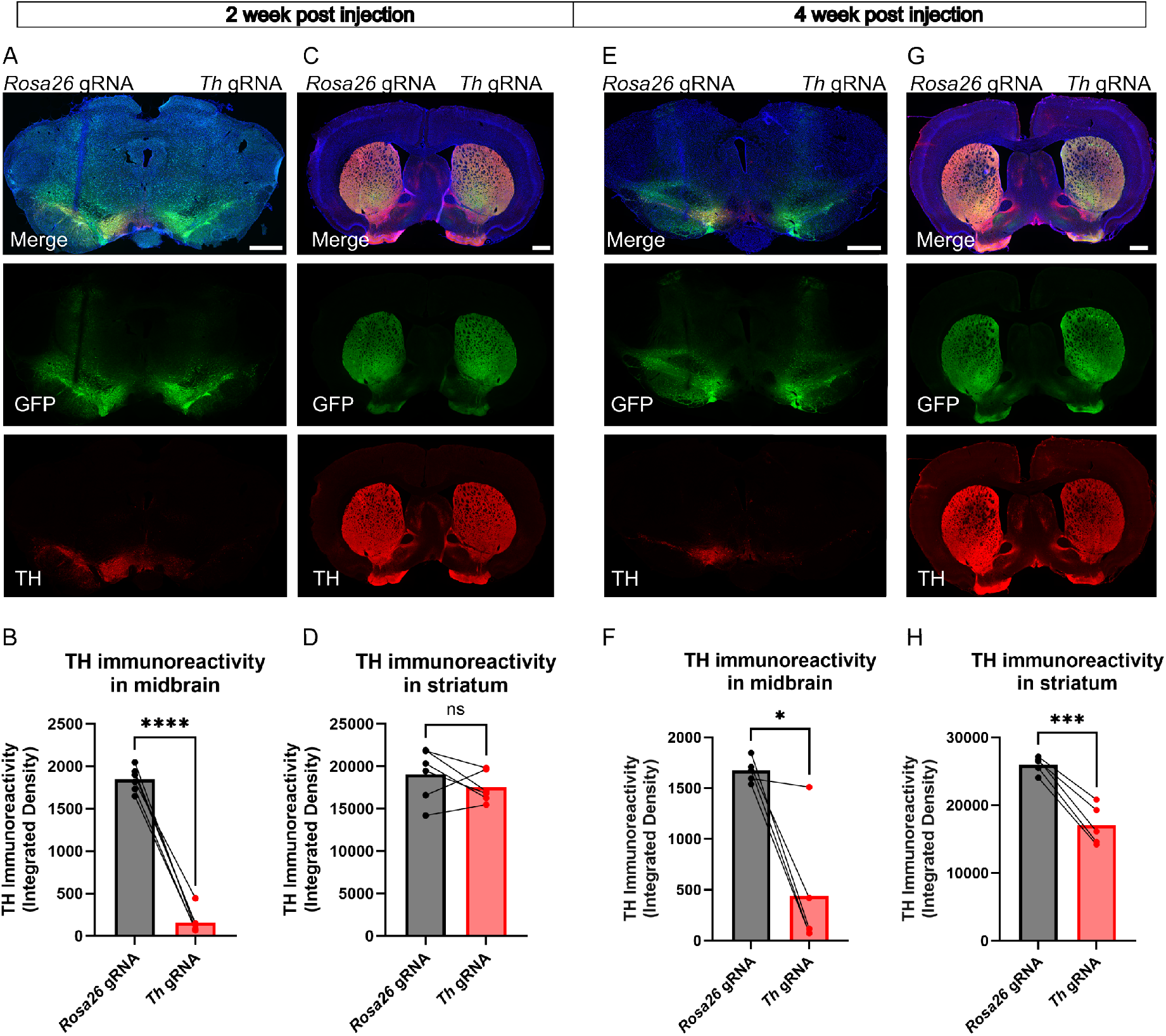
Midbrain and striatum TH knockout two weeks post gRNA injection. (A and C) Representative images of midbrain (A) and striatum (C) 2 weeks after bilateral injections of AAV1-EF1α-eGFP-*Rosa26*gRNAs (left) and AAV1-EF1α-eGFP-*Th* gRNAs (right) into the midbrain of Dat-iCre x LSL-Cas9 rats. GFP fluorescence (green) served as a marker for gRNA delivery. (B, D,F, H) Quantification of TH immunoreactivity was assessed and shown as integrated density. ROIs were drawn using GFP fluorescence. (B and D) Each data point represents 1 animal, where 3 *Rosa26* or *Th* gRNA hemispheres from coronal sections were averaged for each animal (n=6 animals, 3M, 3F). Paired t-tests were performed: B) t=20.38, df=5, ****p<0.0001. D) t=1.219, df=5, p = 0.2772. TH immunoreactivity is significantly reduced in midbrain (B) but not striatum (D) two weeks post gRNA injection. (E and G) Representative images of midbrain (E) and striatum (G) 4 weeks after bilateral injections of AAV1-EF1α-eGFP-*Rosa26*gRNAs (left) and AAV1-EF1α-eGFP-*Th* gRNAs (right) into the midbrain of Dat-iCre x LSL-Cas9 rats. Each data point represents 1 animal, where 3 *Rosa26* or *Th* gRNA hemispheres from coronal sections were averaged for each animal (n=5 animals, 3M, 2F). Pair t-tests were performed: F) t=4.167, df=4, ***p<0.0141. H) t=9.625, df=4, ***p<0.0007. TH immunoreactivity is significantly reduced in midbrain (F) and striatum (H) four weeks post gRNA injection. Scale bars = 1mm

To understand the functional impact of *Th* knockout on dopamine neurons, we performed fast-scan cyclic voltammetry in dorsal striatal brain slices. We observed minimal differences in dopamine (DA) release dynamics between the control and *Th* knockdown striatal hemispheres (Fig. 2). A graphical depiction of dopamine traces (Figure 2A-C) is followed by signal quantification (Figure 2D-E). No significant differences in DA release were observed under baseline conditions (Figure 2D). Under baseline conditions, *Th* gRNA slices had slightly reduced decay times, relative to *Rosa26* gRNA slices (Figure 2A,B). Consistent with this, under baseline conditions, a small but significant increase in the calculated DA uptake parameter (*k_U_*, in s^-1^) was observed in *Th* gRNA slices (Figure 2E). To further assess the release capacity of striatal DA terminals, we measured dopamine release in the presence of the potassium channel blocker 4-aminopyridine (4-AP; 30 µM, Figure 2C - E) which prolongs depolarization of the nerve terminal, thereby enhancing DA release^29–31^. While 4-AP significantly enhanced release in both *Rosa26* gRNA and *Th* gRNA slices, no difference was observed between groups (Figure 2D). Interestingly, 4-AP slightly but significantly reduced DA uptake (see inset, Figure 2C) in both *Rosa26* gRNA and *Th* gRNA slices (Figure 2E). This could either reflect less efficient DA transporter-mediated uptake during depolarization of nerve terminals, or perhaps saturation of uptake at increased DA levels^32^.

**Figure 2.**
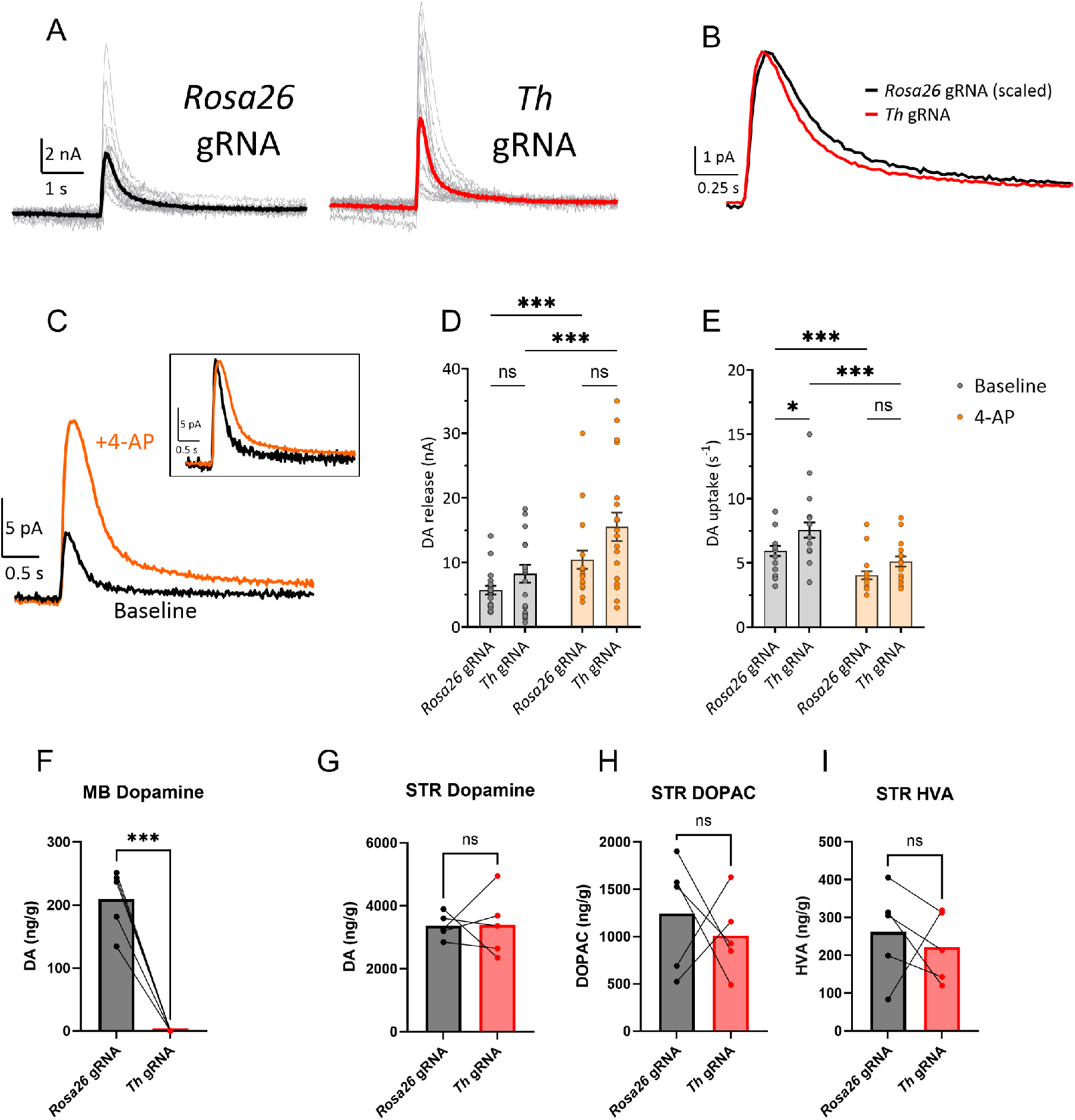
Effect of TH knockdown on dopamine release and uptake. (A) Mean responses are shown for both the *Rosa26* gRNA (black) and *Th* gRNA (red) hemisphere. Individual recordings are shown in light gray. (B) Mean response for the control recordings is scaled to the peak amplitude of the mean *Th* gRNA response. Note the decay of the *Rosa26 gRNA* response is slightly prolonged, relative to the *Th* gRNA response. (C) Effect of 4-AP (30 µM) on the DA signal. Inset shows the baseline response scaled to the peak amplitude of the 4-AP response. (D) DA release under baseline conditions (gray) and in the presence of 4-AP (orange). Under both baseline conditions and in the presence of 4-AP, release did not significantly differ between control and *Th* gRNA (two way RM-ANOVA, F_1,36_ = 3.44, p =0.07; Baseline control vs. baseline TH-, p = 0.68; 4-AP control vs. 4-AP TH-, p = 0.08, Sidak post-hoc). 4-AP significantly increased DA release in both *Rosa26* and *Th* gRNA (two way RM-ANOVA, F_1,36_ = 74.05, p <0.001; ***p <0.001, *Th* gRNA baseline vs. *Th* gRNA 4-AP; *Rosa26* baseline vs. *Rosa26* 4-AP, Sidak-post hoc). (E) DA uptake summary: Under baseline conditions, uptake was slightly but significantly increased in *Th* gRNA slices, relative to control slices (two way RM-ANOVA, F_1,36_ = 5.64, p =0.02; *p=0.04, baseline *Rosa26* gRNA vs. baseline *Th* gRNA, Sidak post-hoc). This increase was absent following 4-AP treatment (4-AP *Rosa26* gRNA vs. 4-AP *Th* gRNA, p =0.30, Sidak post-hoc). Note that 4-AP significantly decreased uptake in both *Rosa26 gRNA* and *Th* gRNA slices (two way RM-ANOVA, F_1,36_ = 76.18, p <0.001; ***p< 0.001, *Rosa26 gRNA* baseline vs. *Rosa26 gRNA* 4-AP; *Th* gRNA baseline vs. *Th* gRNA 4AP, Sidak post-hoc). (F-I) HPLC measuring dopamine and dopamine metabolites from Dat-iCre x Cas9 brain punches two weeks post gRNA injection. (F) DA concentration (ng/g) from midbrain punches (paired t test, t = 9.35, df = 4, ***p = 0.0007). (G) DA concentration (ng/g) from striatal punches (paired t test, t = 0.07, df = 4, p = 0.95). (H) DOPAC concentration (ng/g) from striatal punches (paired t test, t = 0.55, df = 4, p = 0.61). (I) HVA concentration (ng/g) from striatal punches (paired t test, t = 0.55, df = 4, p = 0.61).

To assess the impact of *Th* knockout on total levels of dopamine and dopamine metabolites, we performed high performance liquid chromatography (HPLC) on brain punches from the dorsal striatum and midbrain, 2 weeks following injection with *Th* gRNAs. In the midbrain, there was no detectable dopamine in the *Th* gRNA group, confirming severe deficits in dopamine production (Figure 2F) consistent with the observed reduction in midbrain TH immunoreactivity. In contrast, consistent with no change in striatal TH immunoreactivity, there was no significant reduction in striatal DA, or the metabolites DOPAC or HVA (Figure 2G-I). Given our goal is to identify the early transcriptional signature of disrupted dopamine signaling, we took advantage of the distinct differences in the timeline of TH and dopamine loss between the midbrain and striatum to determine transcriptional changes that occur even before the striatum exhibits these losses.

### *Th* knockout in adult rat midbrain changes transcriptomes of specific neurons and glia

To further understand the impact of TH and dopamine loss on the surrounding cell populations in the midbrain, we performed single nucleus RNA-sequencing on midbrain tissue punches injected with *Rosa26* or *Th* gRNAs. We pooled 4 animals per group, with the *Rosa26* gRNA injected samples stemming from the contralateral hemisphere of the *Th* gRNA injection, so that one hemisphere of each animal was represented in both groups. After quality control read filtering steps, we had ∼10K viable nuclei per experimental group. Using canonical gene markers, we identified 10 unique clusters represented in both datasets (Figure 3A)^33,34^. Six of the clusters represented glial cell types of microglia, astrocytes, and oligodendrocytes throughout differentiation stages: oligodendrocyte progenitor cells (OPC), newly formed oligodendrocytes (NFOL), myelin forming oligodendrocytes (MFOL), and mature oligodendrocytes (MOL). The 4 other clusters were neuronal cell types which included dopaminergic, GABAergic, glutamatergic, and Spp1+ glutamatergic. Spp1+ glutamatergic neurons are a subpopulation of glutamatergic neurons found in the red nucleus, which is a midbrain region dorsal to the ventral tegmental area and substantia nigra (SN) ^35^.

**Figure 3.**
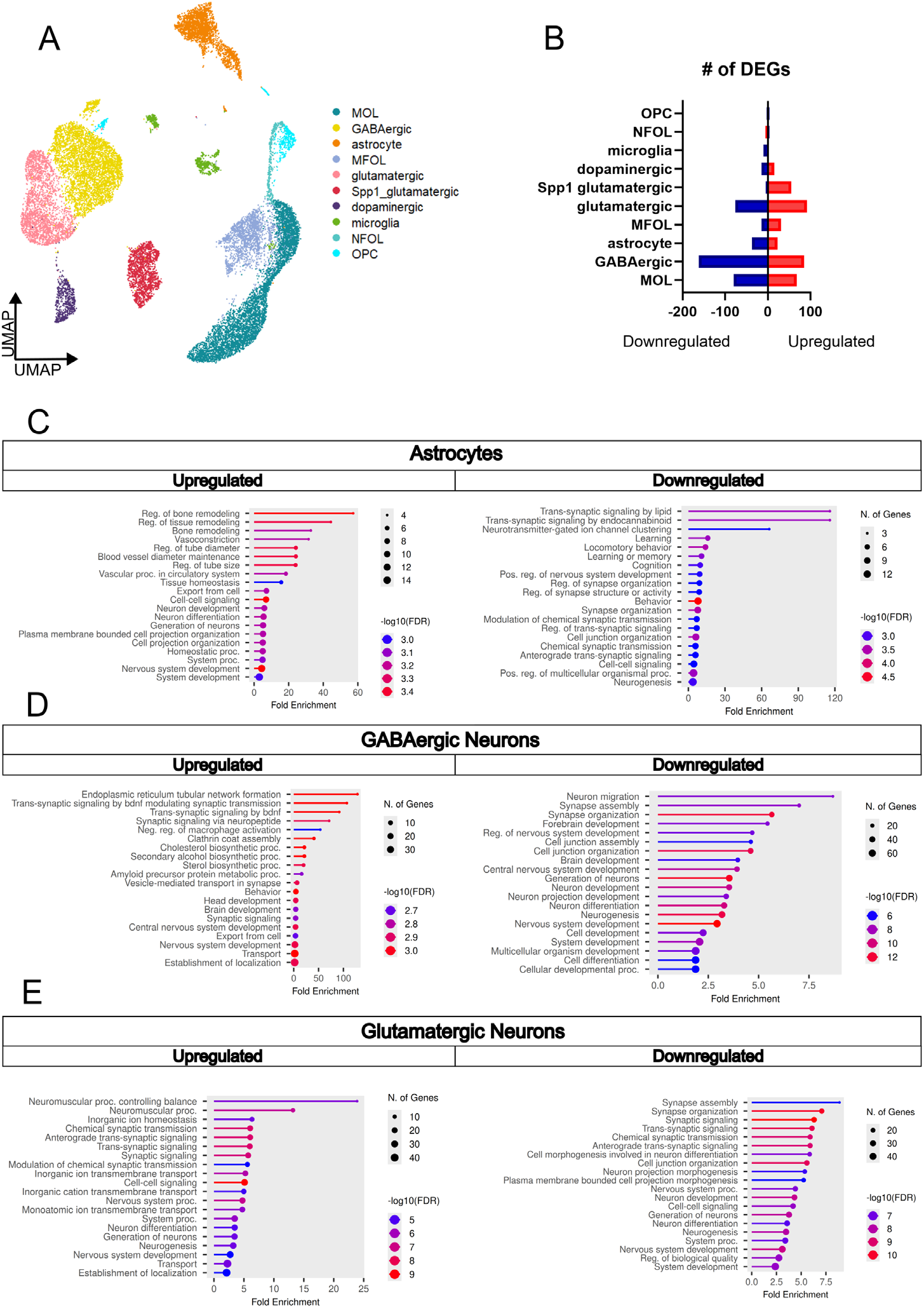
Midbrain snRNAseq after Th knockout shows transcriptional changes in neurons and astrocytes. (A)UMAP representing cell clusters across integrated midbrain dataset. (B) Number of DEGs identified in each cell cluster when comparing *Th* gRNA dataset to *Rosa26* gRNA dataset within clusters. Blue bars on left represent downregulated genes and red bars on right represent upregulated genes. (C – E) Lollipop plots of enriched GO: Biological Processes pathways from significantly differentially expressed genes identified through ShinyGo gene enrichment tool. Color indicates -log_10_(FDR), dot size represents number of DEGs identified in pathway, and x-axis indicates fold enrichment. Cell clusters shown are (C) astrocytes, (D) GABAergic neurons, and (E) glutamatergic neurons.

Next, we performed differential gene expression (DEG) analysis, where we identified differentially expressed genes between *Rosa26* gRNA and *Th* gRNA groups within cell clusters (Figure 3B). Despite having over a 3-fold decrease in *Th* expression, there were only ∼30 total genes significantly changed in the dopaminergic cluster after *Th* knockdown. In the *Rosa26* gRNA group, there are 470 nuclei that constitute the dopaminergic cluster, which represents 4.6% of the total nuclei captured. However, in the *Th* gRNA condition, there are only 181 captured nuclei, representing 1.7% of the total nuclei (Figure S1A). This may suggest that mutating *Th* was lethal for some cells. Among the glial clusters, mature oligodendrocytes and astrocytes showed the largest number of DEGs. Using ShinyGo pathway bioinformatics database, we identified significantly enriched pathways ^36^. Mature oligodendrocytes showed no significantly enriched pathways within their upregulated genes, but there was an enrichment in downregulated genes related to myelination, inflammatory signaling, neuronal development, and neuronal differentiation (Figure S1B). In astrocytes, there was an upregulation of pathways related to vascular regulation, tissue remodeling, neuronal development, and cell to cell communication and a downregulation in pathways related to synaptic transmission, synaptic organization and structural support, neuronal development, cognition, and cell-to-cell communication (Figure 3C).

In GABAergic neurons, genes related to secretory function, neuropeptide and BDNF-related signaling, membrane trafficking and lipid synthesis were increased and genes related to neuronal development and differentiation, synaptic structure, synaptic organization, and brain development were decreased (Figure 3D). These data suggest membrane remodeling in organelle and vesicle machinery, potentially contributing to synapse or dendritic spine remodeling. In glutamatergic neurons, there was an upregulation in pathways related to synaptic signaling, ion homeostasis, developmental/plasticity linking programing and a downregulation in pathways related to synaptic organization, synaptic transmission, developmental/plasticity linking programing. Surprisingly, glutamatergic neurons had the pathway “Transsynaptic signaling” (GO:0099537) enriched in both the up and downregulated pathways. To further explore this bidirectional change, we looked closer at the genes represented. Within the “Transsynaptic signaling” pathway, genes related to general neuron firing such as active zone activity (Snap25, Vamp1), ion handling (Atp2b2, Atp1a3), and axon structure (Nefh, Nefl) were increased. Conversely, genes that decreased were related to more specialized synaptic functions, such as postsynaptic receptors (Grm5, Grik2), adhesion proteins (Ptprd, Cntnap2), and regulators of plasticity (Prkcb, Nos1). Based on these results, TH and dopamine loss in the midbrain leads to an upregulation of genes that promote neuronal activity and a downregulation of genes that contribute to synaptic structure maintenance and synapse specialization.

### Midbrain loss of TH and dopamine leads to cell-specific transcriptomic changes in the striatum

Substantia nigra dopaminergic neurons project to the dorsal striatum, where loss of dopamine signaling causes motor symptoms associated with PD. Although we did not detect changes in striatal dopamine levels, release or TH immunoreactivity, we hypothesized that the early loss of nigral DA production would affect the striatal transcriptome in a way to reflect early adaptations to the loss of dopamine production. In parallel with midbrain samples, we performed single nucleus RNA sequencing on dorsal striatum samples from animals that had received midbrain injections of *Th* and *Rosa26* gRNAs. After filtering the reads for quality control, we had approximately 9.6K and 14.6 K viable nuclei for the *Rosa26* gRNA and *Th* gRNA striatum groups, respectively. Using canonical cell markers, we identified 13 unique cell clusters (Figure 4A)^33,34,37,38^. Six of the clusters represented glial subtypes of microglia, astrocytes, and oligodendrocytes throughout differentiation: OPCs, committed oligodendrocyte progenitors (COP), MFOL, and MOLs. Unlike the midbrain clusters, we did not identify a unique cluster for NFOLs, but instead identified COPs, which exist earlier in oligodendrocyte differentiation. COP clusters express high levels of Neu4, Sox6, Bmp4, and Gpr17 where as NFOLs express those markers at lower levels and have upregulated expression of Tcf7l2 and Casr^34^. The other 7 clusters represent the neuronal clusters identified in the STR: Dopamine receptor 1 (DRD1) medium spiny neurons (MSNs), Dopamine receptor 2 (DRD2) MSNs, carbohydrate sulfotransferase (Chst9) MSNs, Parvalbumin + (PV) interneurons (PVIs), LIM Homeobox 8 + (Lhx8) interneurons, cholinergic interneurons, and somatostatin + (SST) interneurons.

**Figure 4.**
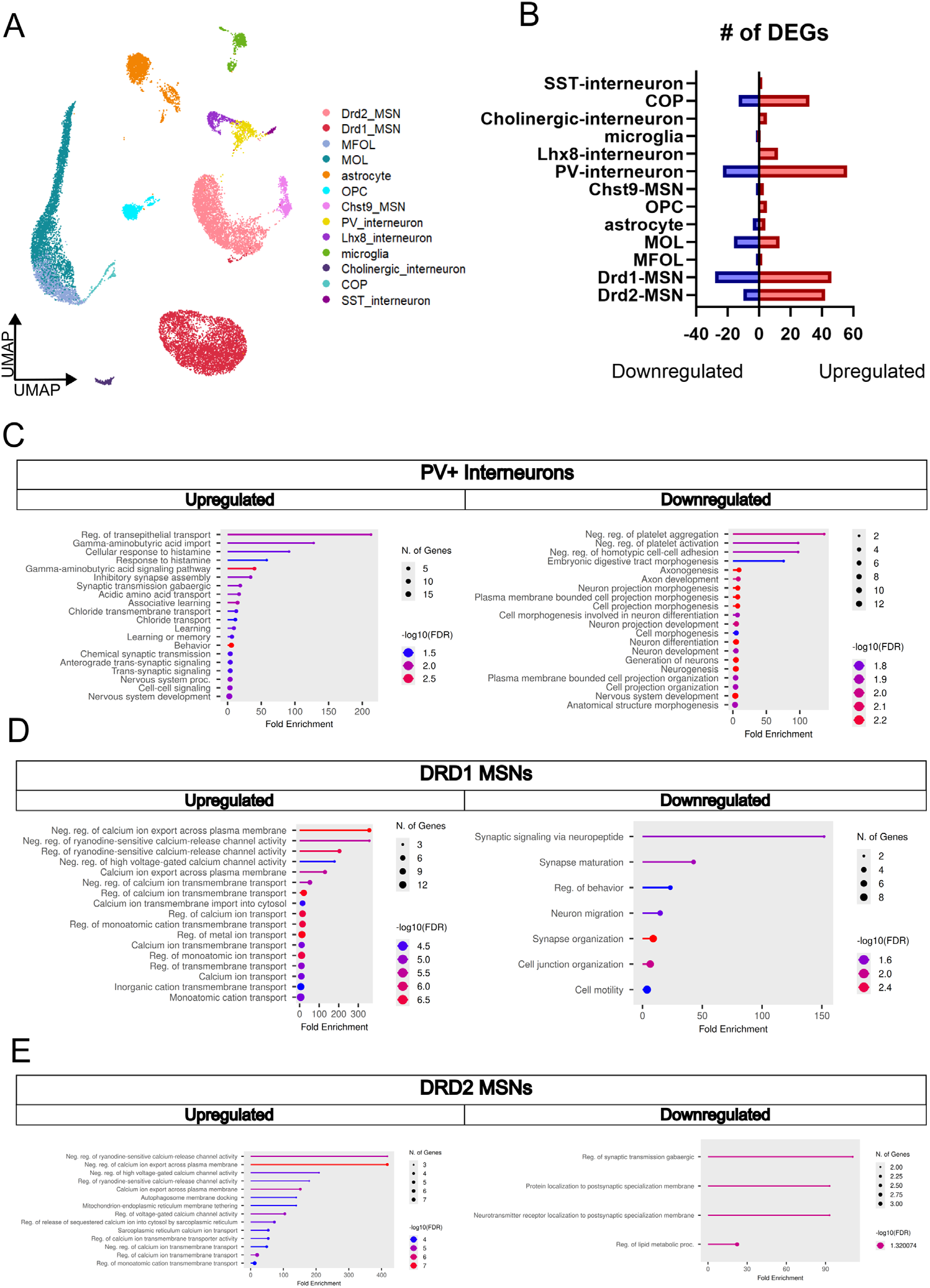
Striatal snRNAseq after midbrain Th knockout shows transcriptional changes in GABAergic neurons. (A)UMAP representing cell clusters across integrated striatum datasets. (B) Number of DEGs identified in each cell cluster when comparing *Th* gRNA dataset to *Rosa26* gRNA dataset within clusters. Blue bars on left represent downregulated genes and red bars on right represent upregulated genes (C-E) Lollipop plots of enriched GO: Biological Processes pathways from significantly differentially expressed genes identified through ShinyGo gene enrichment tool. Color indicates -log_10_(FDR), dot size represents number of DEGs identified in pathway, and x-axis indicates fold enrichment. Cell clusters shown are (C) PV+ interneurons, (D) DRD1 MSNs, and (E) DRD2 MSNs.

Differential gene expression analysis between *Rosa26* gRNA and *Th* gRNA groups identified 5 clusters with greater than 20 differentially expressed genes: COP, MOL, PVIs, DRD1 MSNs, and DRD2 MSNs (Figure 4B). The COP upregulated DEGs include enrichment of pathways related to myelination, glial differentiation, and cell development. COP downregulated genes include enrichment in pathways such as neuron or axon extension, brain development, and plasma membrane organization. It is of note that every downregulated pathway includes Tenascin-R (Tnr) and Semaphorin-5A (Sema5a), both of which were decreased 2.6 fold in the *Th* gRNA group. Tnr and Sema5a are expressed by oligodendrocytes and involved in mediating axonal guidance. Mutations in both Tnr and Sema5a are considered risk factors for familial PD ^39,40^. In mature oligodendrocytes, there was an upregulation of genes related to mRNA splicing and a downregulation of genes related to receptor internalization (Figure S2A). Overall, the transcriptional changes in STR MOLs were less robust than the MB MOL cluster, but both showed a downregulation in receptor mediated endocytosis (Figure S2B). The data suggest that there is a change to oligodendrocyte differentiation and its signal-mediated regulation of neuronal development.

In the neuronal clusters, the 3 cell types that had the highest number of differentially expressed genes were PVIs, DRD1 MSNs and DRD2 MSNs. PV+ interneurons had an upregulation of enriched pathways related to GABAergic function, Chloride homeostasis, cell-to-cell communication, and amino acid signaling (Figure 4C). Genes upregulated within these pathways included GABA transporters (Slc32a1, Slc6a1) and GABA receptors (Gabrb2, Gabra1) indicating an overall shift in GABAergic phenotype within the PVIs. There was a downregulation of enriched pathways related to axon development, neuronal differentiation, neurite remodeling, and adhesion remodeling. DRD1 MSNs had increased expression of genes related to calcium ion homeostasis, RYR-ER calcium release, and ion transport and decreased expression of genes related to synaptic signaling, synaptic organization, and neuronal migration Figure 4D). Similarly, DRD2 MSNs showed an upregulation of calcium ion homeostasis, ion transport, and ER-membrane tethering while demonstrating a downregulation of genes related to localization to the synaptic membrane and lipid metabolism (Figure 4E). Overall, PV+ interneurons, DRD1 and DRD2 MSNs showed increased expression of genes related to ion handling and ion homeostasis, with a focus on calcium homeostasis for the MSNs. These clusters also showed a downregulation of genes related to neuronal development or function, with DRD1 MSNs exhibiting more downregulated genes than the other neuronal clusters.

To assess if striatal transcriptional changes impact electrophysiological properties of MSNs, we performed whole-cell patch clamp electrophysiology on striatal MSNs 2 weeks after *Rosa26* or *Th* gRNA injections. Whole-cell current-clamp recordings revealed no significant differences between *Rosa26* gRNA and *Th* gRNA neurons in resting membrane potential, input resistance, rheobase, action potential half-width, mean interspike interval, adaptation ratio, or afterhyperpolarization amplitude (Figure S3 A-G, supplemental table 1). Analysis of firing frequency-current relationships using REML mixed-effects model demonstrated a significant effect of injected current, but no main effect of treatment and no treatment × current interaction indicating similar excitability across current injection levels in *Rosa26* gRNA and *Th* gRNA neurons (Figure S3 H-I, supplemental table 2). It is likely that the transcriptional changes observed are too early to manifest into measurable physiological changes but may lead to stronger effects over time.

### Neuregulin1 expression is decreased in dopaminergic neurons after *Th* knockout

Despite striatal TH and dopamine remaining unchanged, numerous striatal cell types exhibited transcriptional differences when compared to control suggesting that nigral *Th* knockout alters nigrostriatal signaling that precedes reductions in striatal dopamine. Despite having over a 3-fold decrease in *Th* expression, there were only 29 total genes changed in the nigral dopaminergic neuron cluster after *Th* knockout (Figure 5A). There were no significantly enriched GO Biological Processes pathways. The most upregulated gene was *Mesencephalic astrocyte-derived neurotrophic factor* (Manf), which is an ER stress response protein, known to have trophic effects on DANs and is upregulated in PD^41,42^. Within the dopamine synthesis pathway, only *Th* and *aldehyde dehydrogenase 1A1* (Aldh1a1), which is involved in dopamine breakdown, were altered (downregulated)^43^. The most downregulated gene was *neuregulin1* (*Nrg1*), which exhibited over a 20-fold reduction (log2FoldChange = -4.44). NRG1 is a neurotrophic factor critical for growth and differentiation of neurons and glial cells and is known to play an important role in tuning synaptic strength^44^. With this in mind, we sought to confirm reduction in NRG1 in the midbrain of *Th* gRNA injected animals using immunohistochemistry. Compared to the contralateral *Rosa26* gRNA injected hemisphere, there was a 27% reduction in NRG1 immunoreactivity in the midbrain of *Th* gRNA injected animals (Figure 5B). Based on our data, loss of tyrosine hydroxylase and dopamine in the midbrain leads to a reduction in *neuregulin1* expression that may be an early marker of dopamine dysfunction in Parkinson’s disease.

**Figure 5.**
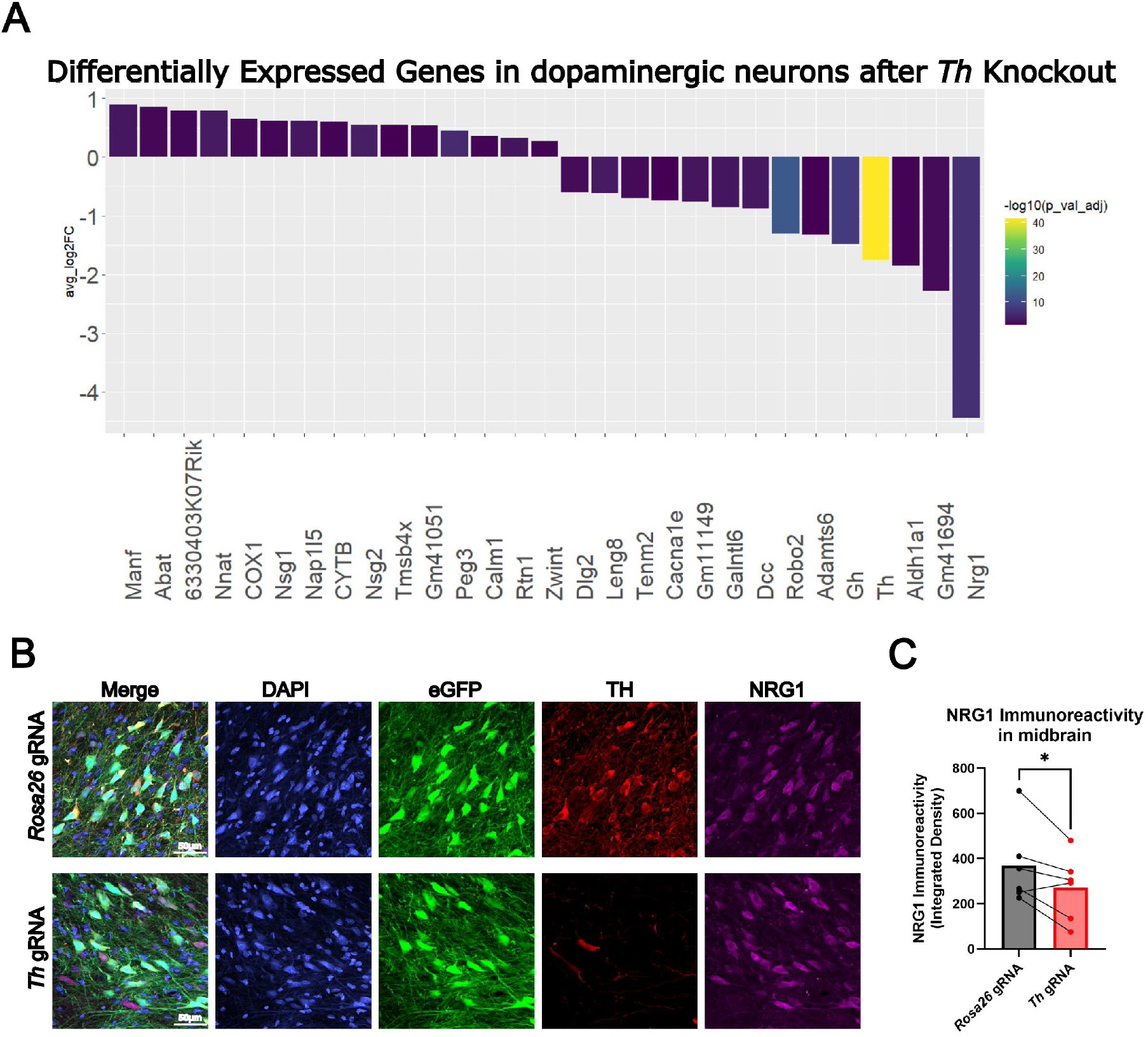
Neuregulin expression is significantly reduced in dopaminergic neurons after TH KO. (A)Significant DEGs (*Th* gRNA vs *Rosa26* gRNA) in midbrain dopaminergic cluster. y-axis is average log2FoldChange and color depicts adjusted p-value. (B) Representative images of midbrain 2 weeks post 2 weeks after bilateral injections of AAV1-EF1α-eGFP-*Rosa26*gRNAs (top) and AAV1-EF1α-eGFP-*Th* gRNAs (bottom). Images include individual channels of GFP (green), TH (red), NRG1 (magenta). Scale bar = 50µm. (C) Quantification of NRG1 immunoreactivity from 20x whole section scans (not shown) of same tissue used in high magnification images in B, shown as integrated density (n=6, 3M, 3F). ROIs were drawn using GFP fluorescence. Each data point represents 1 animal, where 3 *Rosa26* or *Th* gRNA hemispheres from coronal sections were averaged for each animal (n=6 animals, 3M, 3F). Pair t-tests were performed: t=2.610, df=5, *p = 0.048.

## Discussion

Loss of dopamine production in disease leads to debilitating and irreversible motor dysfunction. While nigral dopamine loss occurs at different timepoints within PD, DLB, and MSA, studies suggest that by the time patients being experiencing motor symptoms in PD, they have already lost >30% of their dopaminergic neurons, and the dopaminergic degeneration has been occurring for years prior to diagnosis^6,22,23^. Understanding the early impact of dopamine loss on surrounding cell types in the nigrostriatal pathway may help diagnose synucleinopathies and understand the contribution of these cellular changes to the pre-motor stages of the disease. While other animal models recapitulate aspects of dopamine depletion diseases that occur after dopaminergic cell death, we describe herein a model for studying the early consequences of dopaminergic neurons losing the ability to produce dopamine in adult rat midbrain.

Adult *Th* knockout using Dat-iCre x LSL-Cas9 rats is a promising CRISPR-based approach to study the early transcriptional consequences of disrupting the dopamine synthesis pathway. Injection of *Th* gRNAs is performed unilaterally to create a hemilesion in the nigrostriatal pathway. For nearly 60 years, unilateral lesioning of midbrain dopaminergic neurons with 6-OHDA^45^ and MPTP^46^ has been used to generate hemiparkinsonian rodent models. Hemiparkinsonian models allow for within-subject analysis of dopamine and dopaminergic neuron loss by comparison to the contralateral side. In addition, unilateral lesioning creates internal animal behavioral biases that can be measured through assays like drug-induced rotational bias and elevated body swing test^47–50^. However, neurotoxin models lead to rapid loss of DANs, where loss can begin within hours of injection^14,15^. With *Th* gRNA injections, we cause a gradual loss of dopamine production and signaling, which allows us to study early cellular changes in response to decreased dopamine. Our model also offers temporal control of TH and dopamine loss which allows us to examine cellular transcriptional changes in the adult midbrain that would occur prior to when PD patients exhibit motor symptoms. *Th^-/-^* mice are embryonic lethal unless supplemented with L-DOPA due to TH’s role in synthesizing not only dopamine but norepinephrine and epinephrine as well^51^. Little work has been done with dopaminergic neuron specific *Th* KOs^52,53^. The most common genetic model of dopamine loss is the *Pitx3* ^-/-^ mouse, which is a powerful tool that recapitulate many aspects disrupted dopamine signaling, however these animals are born without a functional dopaminergic system, which does not model the loss of DANs in adulthood, as observed with most forms of idiopathic PD, DLB, and MSA^10,11,20^. In our CRISPR-based model *Th* is knocked out in adulthood, similar to that of human synucleinopathies.

We found that there were detectable transcriptional changes to glia and neurons associated with the midbrain-striatal pathway only weeks after *Th* knockout in dopaminergic neurons. We identified global alterations in synaptic organization and signaling pathways in midbrain and striatal neuronal populations that preceded loss of TH and dopamine in the axon terminals (Figure 6). One of the most notable changes in gene expression after *Th* knockout in dopaminergic neurons was a ∼20-fold reduction in the expression of *neuregulin1,* a strong regulator of neuronal plasticity. This reduction may contribute to transcriptional changes that occur early in PD and other dopamine depletion diseases, in parallel with dopamine loss^44,54^. NRG1 signaling via its receptor, ERBB4, is tightly connected to the dopaminergic system. For example, administration of recombinant NRG1 induces striatal dopamine release and neonatal exposure to recombinant NRG1 increases tyrosine hydroxylase expression ^55,56^. Additionally, systematic administration of NRG1 protects against 6-OHDA induced SNpc degeneration, and in one study, plasma levels of NRG1 were significantly lower in Parkinson’s patients compared to controls, however the decrease does not correlate with clinical severity ^57,58^. Our study found that the >20-fold reduction in *Nrg1* occurs quickly after TH loss and in absence of cell death. Although NRG1 might not be an optimal indicator of gradual dopaminergic neurodegeneration, our data suggests it may be an early biomarker for dopamine loss.

**Figure 6.**
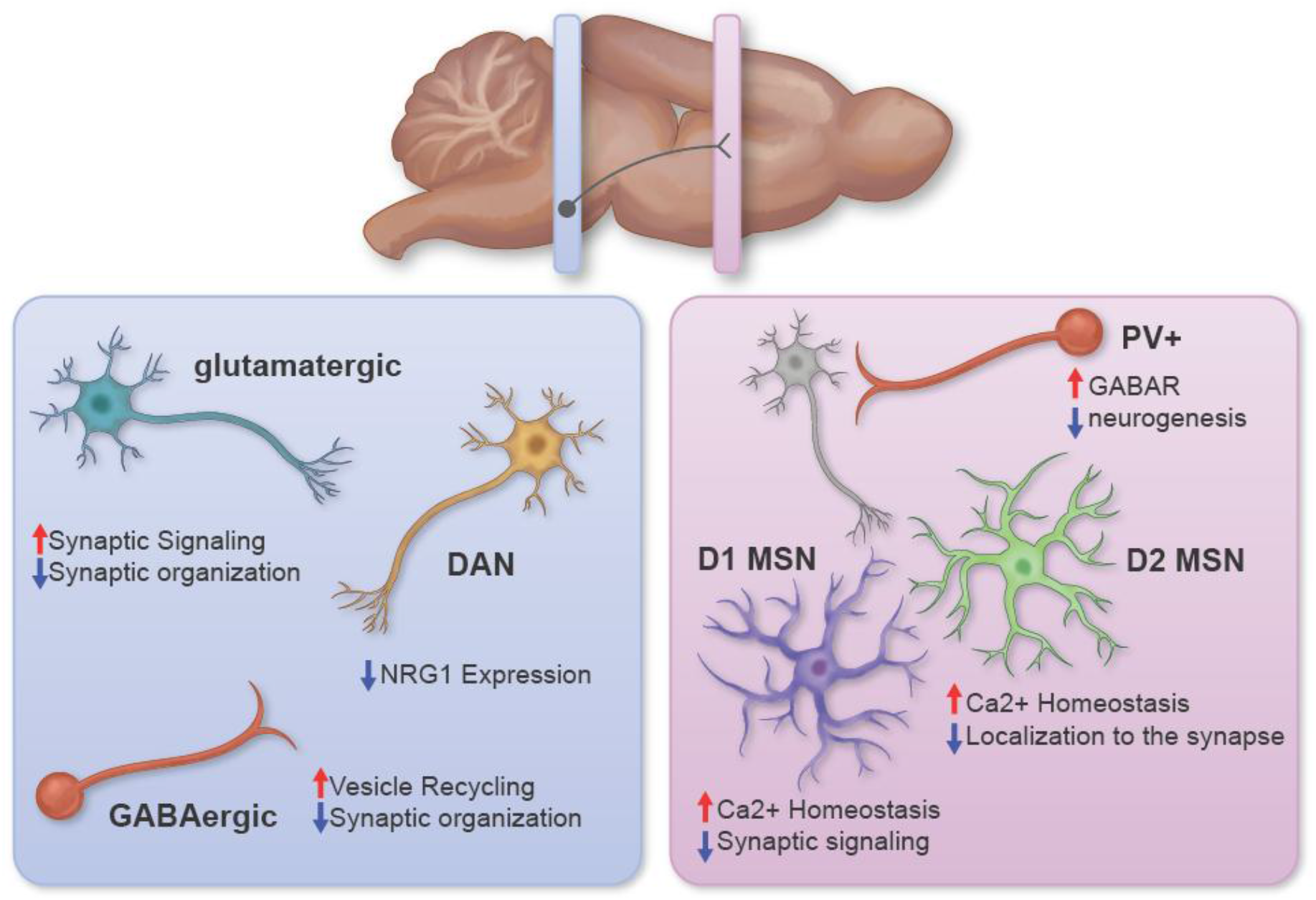
Th knockout causes global alterations to neuronal signaling pathways. Schematic summarizing changes to GO: Biological processes pathways identified in snRNAseq.

We observed mRNA changes in other midbrain neurons as well. Several of these changes might also be related to changes in NRG1 levels. GABAergic neurons exhibited an increased expression of genes enriched in pathways related to synaptic vesicle recycling and remodeling of secretory machinery. They showed a downregulation in genes enriched in synaptic organization. In PD, midbrain GABAergic neurons become more excitable, which is thought to be due to changes in striatal input after loss of dopaminergic signaling to DRD1 and DRD2 medium spiny neurons^59^. However, NRG1 has strong effects on GABAergic neurons throughout development and adulthood because they express high levels of ERBB4. In development, knockdown of ERBB4 impairs spine development in GABAergic neurons and we observed alterations to synaptic remodeling genes in GABAergic neurons ^44^. Similarly, midbrain glutamatergic neurons showed an increased expression in genes related to presynaptic neuronal firing and a downregulation of genes related to postsynaptic organization and plasticity. Midbrain glutamatergic neurons that project to SNpc dopaminergic neurons have decreased release probability and decreased expression of AMPA and NMDA receptor subunits in a LRRK2^G2019S^ mouse model of PD ^60^. In the adult brain, ERBB4 localizes to excitatory synapses and reverses long term potentiation (LTP) on short time scales when activated by NRG1 ^44^. This reversal is thought to be due to AMPAR internalization. Other studies have shown that reversal of glutamatergic LTP in the hippocampus is dependent on NRG1 induced dopamine release ^61^. Collectively, the published data suggest a loss of both NRG1 and dopamine in the midbrain would have profound effects on glutamatergic neurons and could lead to the observed transcriptional alterations in synaptic organization genes.

Two weeks after *Th* knockout in midbrain dopaminergic neurons, we did not observe decreases in striatal tyrosine hydroxylase or dopamine levels, although we see a significant loss of striatal TH at 4 weeks. The delayed loss of striatal TH is likely due to the faster turnover of TH in the soma, compared to more distal neuronal compartments^28^. Consistent with these findings, we did not observe changes in dopamine release or total levels by week two assessed by voltammetry or HPLC, respectively, suggesting that striatal dopamine metabolism is maintained despite loss of midbrain TH and dopamine at this timepoint. However, there were significant transcription changes in PV+ interneurons, DRD1 MSNs, and DRD2 MSNs. In PV+ interneurons, genes associated with GABAergic function were upregulated and those related to neuronal development were downregulated. Under normal conditions, PVIs suppress “anti-movement” beta rhythms and promote gamma oscillations through feedforward inhibition onto MSNs. In PD, the striatum exhibits higher levels of beta oscillations, which is due to a shift toward increased inhibition of PVIs, leading to disinhibition of MSNs ^62,63^. In our study, we observed increased transcription of GABAergic machinery, which would likely promote PVI hypofunction. In support of this, neurotoxin animal models have demonstrated that degeneration of the nigrostriatal pathway leads to disinhibition of the DRD2-MSN indirect pathway, resulting in increased rigidity ^64^. In a 6-OHDA model, MSNs undergo many changes due to the altered dopaminergic and GABAergic input. In response to disinhibition, DRD2 MSNs demonstrate a homeostatic downregulation in intrinsic excitability and rapid decrease in spine density. Conversely, DRD1 MSNs of the direct pathway show an increased intrinsic excitability but reduced total synaptic strength. Both DRD1 and DRD2 MSNS show dendritic atrophy that was not reversed by L-DOPA administration, suggesting that this is not caused by changes to striatal dopamine ^65^. In our dataset, there is an upregulation of calcium handling pathways in both DRD1 and DRD2 MSNs, which precedes a detectable loss of striatal dopamine in our model. We also observed a downregulation of synaptic organization and neuronal migration genes in DRD1 MSNs, which could be contributing to changes in dendritic arborization. Despite the presence of these transcriptional changes in MSNs, we did not observe changes to baseline electrophysiological properties by whole-cell patch clamp. Thus, at this early timepoint the changes have not yet manifested to alter MSN membrane properties, so further analysis at a later timepoint is needed. Interestingly, in DRD2 MSNs, we observed a downregulation of genes related to GABAergic synaptic transmission and receptor localization to the synaptic membrane, including ERBB4. Given that TH and dopamine levels have not changed in the striatum, it’s likely that other mechanisms are driving the transcriptional changes seen in striatal clusters.

Taken together, our data identify NRG1/ERBB4 signaling as a strong candidate to explain the transcriptional changes in the striatum prior to changes to dopamine. NRG1/Erbb4 signaling increases the power of gamma-band oscillations through its promotion of PVI signaling, suggesting that loss of NRG1 might contribute to decreases in PVI inhibitory tone ^66^. *Geng et al*. 2017, showed that a loss of ERBB4 signaling results in hyperactive GABAergic transmission in striatal MSNs, though this study was focused on the ventral striatum and did not differentiate between DRD1 and DRD2 MSNs, so it remains to be seen how loss of NRG1/ERBB4 signaling would impact these MSN subtypes individually. However, the downregulation of Erbb4 in DRD2 MSNs in our data points to NRG1/ERBB4 signaling involvement. Consistent with this, NRG1/ERBB4 signaling loss in GABAergic neurons impairs dendritic arborization, potentially explaining the dopamine-independent deficits in dendritic arborization observed in 6-OHDA models that may already be detectable through the downregulation in membrane remodeling and synaptic organization pathways seen in MSNs.

Previous single-cell RNA sequencing studies in PD models and human post-mortem tissue have implicated “pan glial” activation in PD progression ^67–70^. We did not observe any pro-inflammatory phenotypes in midbrain glial clusters. While we did observe significant changes to midbrain astrocytes, they were enriched in remodeling and structural support pathways, suggesting that acute loss of midbrain TH is not sufficient to induce pro-inflammatory phenotypes observed in PD. In mature oligodendrocytes in the midbrain, there was a significant downregulation in genes related to myelination and neuronal signaling consistent with observations of decreased white matter in the brains of PD patient and suggests that demyelination may be mediated by loss of dopamine ^71^. The parkinsonian form of MSA exhibits α-synuclein inclusions in oligodendrocytes in the nigra with toxicity of MOLs and demyelination^72,73^. It is possible that nigral degeneration and the consequent loss of dopamine may be contributing to this phenotype. In the striatum, MOLs and COPs also exhibited changes. COPs showed a downregulation of *Tnr* and *Sema5a*, which are involved in cell adhesion and mediating axon guidance. Mutations in both genes have been implicated in PD^39,40^. Interestingly, NRG1 signaling has been implicated in transcriptional changes that occur in astrocytes and oligodendrocytes in Parkinson’s disease^74^. Under normal conditions, NRG1 promotes differentiation, myelination, and survival of oligodendrocytes^44^. In contrast, animals haploinsufficient for NRG1 are hypomyelinated^75^. Due to the central role that oligodendrocytes play in MSA, it has been speculated that NRG1 may be important for disease pathology and therapeutics^8^. In the cases of both midbrain glia and neurons, NRG1 and dopamine signaling are involved in regulation of synaptic strength and neuronal signaling pathways. The transcriptional changes observed in our study support a phenotype of altered neuronal synaptic strength, synaptic organization and glia-mediated neuronal maintenance, which are consistent with changes observed in PD models and may involve NRG1 signaling.

Synucleinopathies and forms of PD are not the only conditions that exhibit dopamine depletion. Substance use disorders such as chronic use of cocaine, methamphetamines, or nicotine, have demonstrated dopaminergic hypofunction overtime^12,76–78^. Dopaminergic reward signaling relies heavily on the VTA which projects to the nucleus accumbens, amygdala, cortex, hippocampus, ventral pallidum, periaqueductal grey, bed nucleus of the stria terminalis, olfactory tubercle, and locus coeruleus^79^. Our model shows that injection of *Th* gRNAs into the midbrain reduces TH immunoreactivity in both the SNpc and VTA. The midbrain punches we collected included VTA cells and SNpc cells, but insufficient markers to separate VTA and nigral dopaminergic neurons and sample size led us to keep them combined in one dopaminergic cluster. Our striatal samples were from the dorsal striatum, which excludes the nucleus accumbens. It is likely that loss of dopaminergic signaling has strong impacts on reward circuitry, but this study focuses on the nigrostriatal pathway. Our model would be useful to study impacts of dopaminergic hypofunction on mesolimbic reward system, but it is beyond the scope of our current study.

We describe a rat model of dopamine loss in the adult midbrain that is unilateral and temporally controlled. The loss of TH occurs fast enough to be a viable experimental model (2-4 weeks), but slowly enough to reveal transcriptional changes in associated cells of relevant brain regions. Our model represents a platform to study the early cellular changes that occur after a loss of dopamine synthesis, but prior to the onset of axonal/terminal DA depletion. Herein we identify a loss of NRG1 in dopaminergic neurons. To the best of our knowledge, our study is the first demonstration that knocking out *Th* is sufficient to decrease NRG1 expression. The robust reduction in NRG1 opens the door for earlier interventions in dopamine depletion diseases like synucleinopathies and identifies neuregulin1 as a potential mediator of early transcriptional changes occurring in surrounding midbrain and striatal cell types that correspond with known disease pathology.

## Materials and Methods

### AAV Vectors

Animals were injected with pAAV Rosa26 gRNA A+B EF1a EGFP (Addgene 113156) and pAAV TH gRNA A+B pair EF1a EGFP (Addgene 113155) previously described in *Bӓck et al* 2019 at a viral titer of 1.95e12 viral genomes/mL. All vectors were produced using the triple transfection method as previously described^42^. All vectors were produced using serotype 1 capsid proteins and titered using digital PCR.

### Animals

All animal experiments were conducted on adult male and female Long Evans Dat-iCre x LSL-Cas9 rats following National Institute of Health (NIH) guidelines for animal research. All experiments were approved were approved by the Animal Care and Use Committees (ACUC) of the Intramural Research Programs at the National Institute on Drug Abuse (NIDA). The rats were group-housed in a 12h light-dark cycle with *ad libitum* access to rodent chow and water. Animals were selected for experiments based on genotyping results. Animals were procured from the NIDA Intramural Research Program’s transgenic breeding program in accordance with approved protocols.

### Intracranial Injections

Adult Long Evans Dat-iCre x LSL-Cas9 rats (2-4 months) were anesthetized in an induction chamber with 4% isoflurane until they were in plane 2 of anesthesia and did not exhibit a withdrawal reflex. Surgical area was shaved before animals were placed in stereotaxic frame (Stoelting Co., Wood Dale, IL). Rats received bilateral intracranial AAV injections into the substantia nigra (SN) at coordinates : AP -5; ML ± 1.8; DV -8 mm from bregma. A total of 1.0µL of AAV was injected into each hemisphere at a rate of 0.5 µL/min. Injections were performed using a Nanofil 10 µL syringe with a 33G blunt needle and UMP4 microinjector pump (World Precision Instruments, Sarasota, FL). Follow the injection, needle was kept in place for an additional 2 min to reduce backflow of AAV.

### Euthanasia and tissue processing for immunohistochemistry

Rats were deeply anesthetized with isoflurane and transcardially perfused with heparinized phosphate-buffered saline (PBS), followed by 4% paraformaldehyde (PFA) solution. Brains were extracted, post fixed in 4% PFA for 2 hrs at 4°C, then incubated at 18% sucrose followed by 30% sucrose before freezing on dry ice and storing at -80°C until use. Brains were coronally sectioned in cryostat at -17°C. The striatum was sectioned at a thickness of 50µm and the midbrain was sectioned at a thickness of 30µm. Tissue was stored in 1x phosphate buffer (PB) with 20% glycerol and 2% DMSO at -80°C.

### Immunohistochemistry

Tissue sections were thawed and washed in PB for 30 min at room temperature (RT). Sections were blocked in PB, 0.3% TritonX-100, and 4% goat serum for 1 hour at RT. Tissue was incubated overnight at 4⁰C in primary antibody and blocking solution. We used Mouse anti Tyrosine hydroxylase (1:1000 Millipore MAB318) and Rabbit anti neuregulin (1:250 ThermoFisher PA5-78627). The next day, tissue was washed for 30 min at RT in PB and incubated in secondary antibody (1:2000) and blocking solution for 2 hrs at RT. Sections were washed with PB for 10 min, stained with DAPI (1:3000) for 10 min at RT, and washed for 10 min in PB before mounting in mowiol mounting reagent

### Microscopy

We imaged whole sections using the Olympus slide scanner (Olympus VS200 slide scanner, Evident Scientific, Waltham, MA). 20x scans underwent 8×8 binning during file conversion to TIFFs. High magnification confocal images were acquired using Nikon system (Nikon Eclipse Ti2 microscope system, NIS-Elements imaging software version 5.42; Nikon Instruments Inc., Melville, NY)

### Densitometry Analysis

Densitometry analysis of coronal midbrain and striatal sections was performed in Fiji/ImageJ (Schindelin et al., 2012) (version 2, NIH). After background subtraction, manually drawn regions of interest (ROI) were drawn around the VTA and SN in midbrain or dorsal striatum in STR of both *Th* gRNA injected and contralateral *Rosa26* gRNA control side based on GFP fluorescence as an injection marker. We measured integrated density of TH or NRG1 immunofluorescence. Measurements from 3 sections were averaged for each animal with an n=3 for each sex (n=6 total).

### Fast Scanning Cyclic Voltammetry

Approximately 2 weeks following *Th* gRNA or *Rosa26* gRNA injections, rats (2 males, 4 females) were anesthetized with isoflurane and decapitated, and striatal brain slices (280 µm) were prepared as previously described^27^. FSCV recordings were performed according to previously published protocols^27,80^. Signals were acquired using HDCV software^32^ and a voltammetric amplifier (EVA-8, Heka Electronik, Germany). Carbon fiber electrodes were positioned at a depth of approximately 150 µm in the slice, and a bipolar stimulating electrode was positioned within ∼100 µm of the recording electrode. Electrical stimulation was delivered to the dorsal striatum using a 1 ms, 150µA constant current pulse. For each subject, 3-4 slices were obtained from each hemisphere (*Rosa26* gRNA and *Th* gRNA), and the same recording electrodes were utilized in both the *Rosa26* and *Th* gRNA slices within each subject. The presence of GFP expression was confirmed prior to each experiment using a stereomicroscope equipped with a fluorescence adapter (Nightsea, Hatfield PA). One animal was removed from analysis due to lack of GFP expression. At each recording location, 3-5 signals were obtained to generate a baseline average response prior to any pharmacological treatment. 4-aminopyridine (30 µM) was applied for 8-10 minutes and 3-5 responses were averaged. Signals were analyzed using a nonlinear curve fitting restricted diffusion model that evaluates DA release and uptake in brain slices ^80^.

### Electrophysiology

Approximately 2 weeks following TH gRNA or control injections, rats (2 males, 3 females) were anesthetized with isoflurane and decapitated, and striatal brain slices (280 µm) were prepared as previously described^27^. For recording, a single brain slice was submerged in a recording chamber (∼300 µL) integrated into a fixed stage of an upright microscope (Olympus BX51WI, Evident Scientific, Needham MA) equipped with differential interference contrast and infrared optics (DIC-IR) and epifluorescence. The presence of GFP expression was confirmed prior to each experiment using fluorescence microscopy. Slices were perfused at a rate of 2-3 ml/min with oxygenated aCSF (in mM: NaCl, 126; KCl, 3; NaH2PO4, 1.2; NaHCO3, 26; Glucose, 11; MgCl2, 1.5; CaCl2, 2.4; using a peristaltic pump (Cole-Parmer, Vernon Hills, IL) and maintained at 32°C using an inline solution heater (Warner Instruments). All recordings were performed in dorsal striatum. Recordings were performed using borosilicate pipette glass (Sutter Instruments, Novato, CA, 1.5 mm O.D. × 0.86 mm i.d.) and a horizontal puller (P-97; Sutter Instruments). Electrodes were filled with a potassium-based internal solution consisting of (in mM): K-gluconate, 140; KCl, 5; HEPES, 10; EGTA, 0.2; MgCl2, 2; Mg-ATP, 4; Na2-GTP, 0.3; Na2-phosphocreatine, 10), ∼ 295–305 mOsm, neutralized to a pH of 7.2 using potassium hydroxide. Electrode resistances were 4–7MΩ when filled with this solution. Whole-cell patch clamp recordings were performed using an Axon Instruments 700B Multi-Clamp amplifier (Molecular Devices, San Jose, CA) and WinLTP software (WinLTP, Bristol, UK). Analyses were performed offline using custom written scripts in MatLab (Mathworks, Natick MA).

### Sample processing for single nucleus RNA sequencing and HPLC

Rats were lightly anesthetized with isoflurane then rapidly decapitated. Brains were removed within 2 minutes and flash frozen in prechilled isopentane for 30 seconds. Brains were coronally sectioned on cryostat at -20°C until reaching approximate anterior/posterior coordinates of 1.5mm from bregma. 1.5mm x 2mm punches of dorsal striatum were collected from each hemisphere. Brains were then sectioned further to -4.5mm from bregma and 1.0mm x 2mm punches of the substantia nigra midbrain region of each hemisphere were collected. Brain punches were frozen at -80°C until used for HPLC or nuclei extraction.

### Dopamine HPLC

Tissue samples for HPLC analysis were thawed on ice and homogenized into 300 ul of aqueous antioxidant solution containing 171 mM perchloric acid, 14 mM acetic acid, 0,43 mM L-cysteine and 0,14 mM oxalic acid. Homogenization was performed by using ultrasound sonicator (GM35-400, Rinco Ultrasonic, Switzerland) continuously for at least 60s. 300 ul of homogenized samples was transferred to 0.5 mL Vivaspin centrifugal filter concentrators (10 kDa MWCO, PES membrane; Sartorius Lab Instruments GmbH & Co. KG, Göttingen, Germany) and centrifuged at 8900 x g at 4C for 35 min. 100 ul of flittered sample was injected into HPLC system for analysis.

The HPLC system for determination of the concentration of neurotransmitters and their metabolites consisted of a solvent delivery pump (Jasco model PU-2080 HPLC Pump, Jasco International Co, Japan), an analytical column (Kinetex C-18 150 mm × 4.6 mm, 2.6 µM particle size; Phenomenex, CA, USA) protected by a 0.5-mm inlet filter and thermostated by a column heater (CROCO-CIL, Cluzeau Info-Labo, France), and a electrochemical detector (ESA Coularray multi-electrode detector system, model 5600A detector and three flow cells (Model 6210, ESA, USA). Each cell consists of four analytical cells (model 5014B cells; Esa InC., Chelmsford, MA, USA). The potentials of analytical cells were set from 30 mV to 360 mV with 30 mV increments. The mobile phase consisted of 0.1 M NaH_2_PO_4_, 7 % v/v MeOH, 450 mg/l EDTA and 450mg/l sodium 1-octanesulfonate, pH adjusted to 3. Flow-rate was set to 1.1 ml/min and column temperature to 45C. 100 ul of the sample filtrate was injected onto the column with a refrigerated autoinjector (Shimadzu SIL-20AC, Shimadzu Corp, Japan). The chromatograms were processed by ESA CoulArray chromatography data system software (Chelmsford, MA, USA). 6 standards from 2 ng/ml to 64 ng/ml for each analyte were prepared to homogenization solution and run before and after the samples together with blank sample containing only homogenization solution.

### Nuclei Extraction and Fluorescence activated nuclei sorting (FANs)

Nuclei were extracted using established protocols (Matson et al 2018) with some modifications as described below. Briefly, we prechilled all buffers and Dounce homogenizers then performed mechanical lysis in lysis buffer (0.32 M sucrose, 10 mM, HEPES pH 8.0, 5 mM CaCl2, 3 mM MgAc, 0.1 mM ETDA, 1 mM DTT, 0.1% Triton X-100). We diluted lysates in prechilled low sucrose buffer (0.32 M sucrose, 10 mM, HEPES pH 8.0, 5 mM CaCl2, 3 mM MgAc, 0.1 mM ETDA, 1 mM DTT) and filtered sample through a 40µm strainer. Samples were centrifuged at 3200 g for 10 min at 4°. After centrifugation, we poured off supernatant and resuspended the pellet in 0.5mL resuspension buffer (1x PBS, 0.4mg/mL BSA, 0.2 U/µL RNAse Inhibitor). Samples were passed through 40µm cell strainer into 3mL cold resuspension buffer. Samples were then centrifuged at 250 g for 10 min at 4°. 2.5 mLs were aspirated to leave 1mL nuclei – resuspension buffer mixture. Samples were then incubated in Vybrant DyeCycle Green nuclear dye (ThermoFisher cat# V35004 1:500) for 15 min on ice.

We used a BD FACS Meldoy sorter equipped with 3 excitation lasers (488nm, 561 nm, and 640 nm) and 9 emission filters for FANs. Nuclei formed a distinct cluster separate from debris and was gated in forward scatter area vs. side scatter area view. We determined intact nuclei by gating for positive vibrant green nuclear signal where we excited samples with 488nm laser and collected emitted fluorescence from 507 to 527nm with a detection PMT voltage for 218. We aimed to sort 15000 nuclei from each of the 4 brain punches for each treatment group. We sorted nuclei from rosa gRNA and Th gRNA samples into separate tubes filled with sample buffer and RNAase inhibitor.

### Nuclei Capture and Sequencing Library preparation

We used the BD Rhapsody Single cell analysis platform for nuclei capture and library preparation. Briefly, nuclei were loaded into BD Rhapsody cartridge (v2) and incubated with barcoded capture beads followed by lysis. We followed the manufacturers protocol: *Single Cell Capture and cDNA synthesis with the BD Rhapsody Single-Cell Analysis System* (DOCID210966) with a few modifications for nuclei capture rather than whole cell capture. Rosa gRNA and Th gRNA samples were run in parallel throughout this protocol. We performed cell counting using Vybrant DyeCycle Green nuclear dye (ThermoFisher cat# V35004 1:500). We added 0.2 U/µL RNAse inhibitor (ThermoFisher cat# AM2682) to Sample buffer for cartridge and bead loading steps. We added 50 µL of 800 U/µL Proteinase K (Machery-Nagel cat# 740506) to lysis buffer. Samples containing barcoded beads bound to polyadenylated nucleic acids were collected. We next performed reverse transcription and exonuclease reactions. For library preparation steps, we followed *BD Rhapsody mRNA Whole Transcriptome Analysis (WTA) and Sample Tag Library Preparation Protocol* (DOC ID 2321712-00). We did not perform any sample tagging, so these steps were excluded. Briefly, we performed random priming and extension, then amplified and purified products using AmpureX magnetic beads. We determined size and quality of products on Agilent 2100 Bioanalyzer. Finally, we indexed *Rosa26* gRNA and *Th* gRNA samples, purified, and checked quality using Agilent 2100 Bioanalyzer.

### Single nucleus RNA sequencing and genome alignment

Samples were pooled for sequencing on Illumina sequences. We sequenced whole transcriptome libraries at ∼60000 reads/nucleus on an Illumina NovaseqX (PE 75bp) through collaboration with the NIH intramural sequencing center (NISC). We followed the BD Rhapsody WTA analysis pipeline on Seven Bridges Genomic platform (v1.11 rev 8) to demultiplex raw sequencing reads and align to rat reference genome (Rn7.2). We generated filtered count matrices with recursive substitute error correction (RSEC) based on the mm10 mouse genome annotation to RN7.2 rat genome^81^ . Raw and filtered sequencing data were uploaded to Gene Expression Omnibus (GEO) in library: GSE341879 (https://www.ncbi.nlm.nih.gov/geo/query/acc.cgi?acc=GSE341879).

### Sequencing Analysis

All analysis scripts are available in our Github repository (https://github.com/NIDA-IRP/TH_KO-snRNAseq). Rhapsody RSEC count matrices were analyzed using Seurat (v5.2.1) in R (4.4.3)^82^. We created Seurat objects from count matrices. Experimental metadata for each nucleus was assigned according to its cartridge. Datasets were filtered for ambient RNA using Soup X and doublets using DoubletFinder ^83,84^. After filtering, the *Rosa26* gRNA and *Th* gRNA datasets were integrated. We applied the MapMyCells function from the Allen Brain atlas to assist in identifying cell types (MapMyCells RRID:SCR_024672). These identifications were added to the metadata. We normalized and scaled data using the 2000 most variable features before applying PCA dimensional reduction using dimensions 1-15 and a resolution of 2 to form clusters. Clusters with low RNA features and RNA counts were removed and data was reclustered^85^. We manually identified cell clusters using canonical markers: Mature oligodendrocytes (Klk6, Trf, Apod, Mal), myelin forming oligodendrocyte (Low: Klk6, High: Mal, Mog, Plp1, Opalin), oligodendrocyte progenitor cell (Pdgfra, Cspg4), committed oligodendrocyte progenitor/newly formed oligodendrocyte (Casr, Neu4, Sox6), Astrocyte (Gfap, S100b, Slc7a10), microglia (Cx3cr1, P2ry12, Tmem119), GABAergic (Slc32a1, Gad1, Gad2), glutamatergic (Slc17a6, Gls), Spp1_glutamtergic (Spp1, Gls), dopaminergic (Slc6a3, Th, Slc18a2), endothelial cell (slc22a8, Esam). Manual cluster identification was cross checked with Allen Brain atlas classifications to confirm assessment. We subsetted data to remove endothelial cells from further analysis. For differential expression analysis, we used the “FindMarkers” package in Seurat using Wilcoxon rank sum test to compare individual clusters between experimental group using false discovery rate corrected p-values. All DEGs for each comparison are listed in SupplementalTable_MB_DEG (MB) and SupplementalTable_STR_DEG (STR). While we pooled both male and female animals in accordance with sex as a biological variable policy, nuclei from individual samples were not labeled so we could not compare differential gene expression by sex.

### Quantification and Statistical Analysis

All results are shown as mean ± standard deviation (SD). Statistical analysis was performed using GraphPad Prism software (GraphPad, San Diego, CA). For comparison between 2 groups, student’s paired t-test was applied. For measurements with 2 factors as in the fast scan cyclic voltammetry measurements, a two-way ANOVA was performed with uncorrected Fisher’s LSD post-hoc test. A p value of <0.05 was considered significant.

## Funding Declaration and Acknowledgements

This research was supported by the Intramural Research Program of the National Institutes of Health (NIH). The contributions of the NIH author(s) were made as part of their official duties as NIH federal employees, are in compliance with agency policy requirements, and are considered Works of the United States Government. However, the findings and conclusions presented in this paper are those of the author(s) and do not necessarily reflect the views of the NIH or the U.S. Department of Health and Human Services. Specifically, this work was supported by the Intramural Research Programs at the National Institute on Drug Abuse. MA and AJ were supported by the Sigrid Jusélius Foundation. We thank Doug Howard and the Genetic Engineering and Viral Vector Core for AAV packaging, the NIH Intramural Sequencing Core for sequencing snRNAseq libraries, and the NIDA Histology Core for use of the VS200 slide scanner. We would like to thank Lauren Brick of NIH Visual Media Services (Baltimore, MD) for assistance with preparing Figure 6.

## Author Contributions

L.G., and B.H. designed experiments. L.G. performed experiments with the assistance of A.E.H. (Figure 1 and 5), A.J. (Figure 2) and A.F.H. (Figure 2 and Figure S3). L.G., A.F.H., and B.H. analyzed data. L.G. and B.H. wrote the manuscript. All authors contributed to reviewing results and writing the manuscript.

## Supporting information

SupplementaryTable_MB_DEG

SupplementaryTable_STR_DEG

---

**Supplemental Figure 1.**
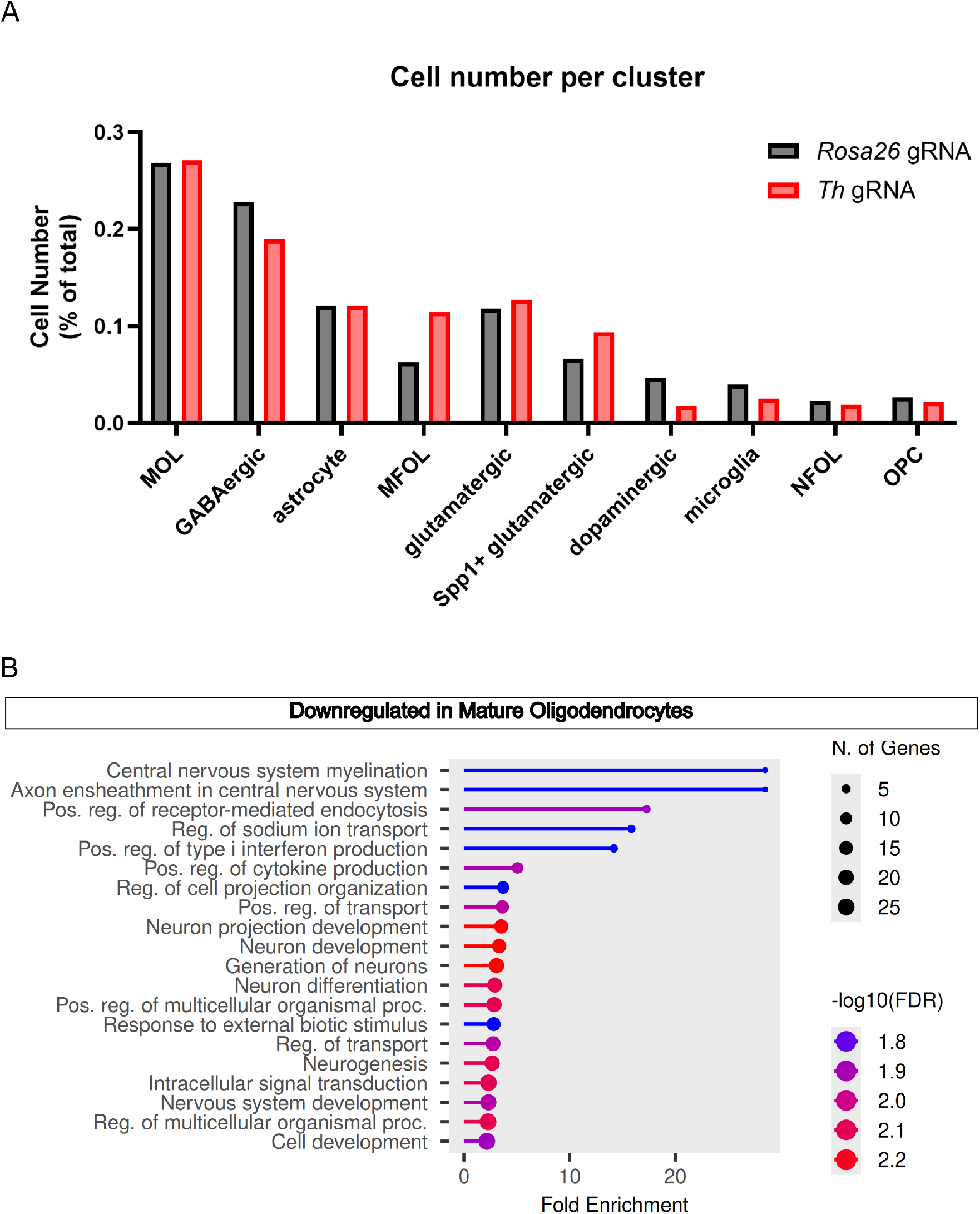
Relative cell numbers per cluster and downregulated pathways inn mature oligodendrocytes from midbrain snRNAseq after *Th* knockout. (A)Relative cell numbers within each cell cluster from midbrain snRNAseq *Rosa26* gRNA (Black) and *Th* gRNA (red) datasets presented as percentage of total cells in dataset (B) Lollipop plots of enriched GO: Biological Processes pathways from significantly differentially expressed downregulated genes in mature oligodendrocyte midbrain clusters. Pathways identified through ShinyGo gene enrichment tool. Color indicates -log_10_(FDR), dot size represents number of DEGs identified in pathway, and x-axis indicates fold enrichment.

**Supplemental Figure 2.**
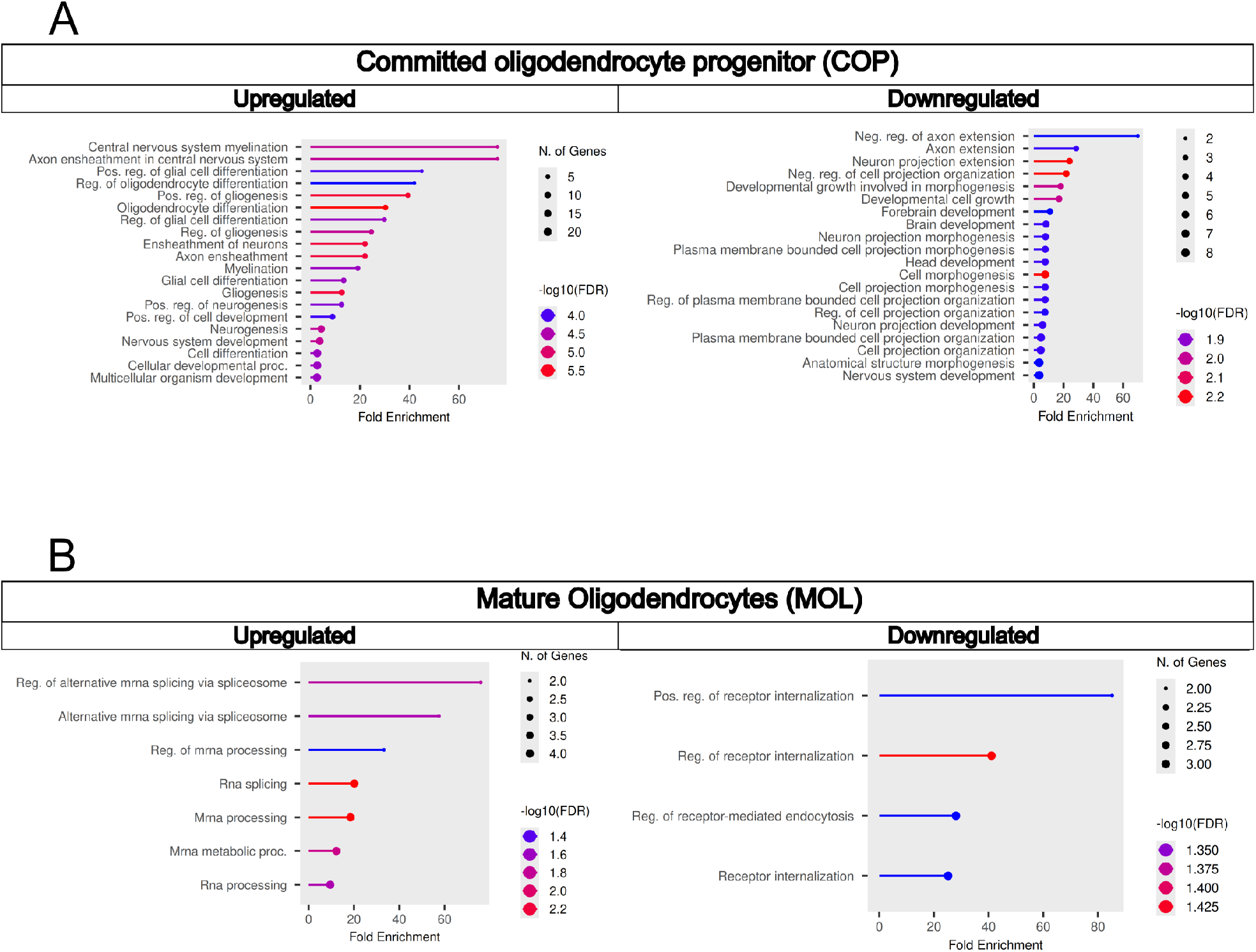
Striatal snRNAseq after midbrain *Th* knockout shows transcriptional changes in oligodendrocytes. (A) Lollipop plots of enriched GO: Biological Processes pathways from significantly differentially expressed genes in committed oligodendrocyte progenitor (A) and mature oligodendrocytes (B) clusters from striatal snRNAseq clusters. Pathways identified through ShinyGo gene enrichment tool. Color indicates -log_10_(FDR), dot size represents number of DEGs identified in pathway, and x-axis indicates fold enrichment.

**Table 1.** Statistical summary of passive membrane properties in MSNs.

| Parameter | Control (n=14)<br>Mean ± SEM (95% CI) | TH KO (n=12) Mean ± SEM (95% CI) | Welch's t(df) | p |
| --- | --- | --- | --- | --- |
| Resting membrane potential (mV) | -75.38 ± 1.70 (-79.05, -71.72) | -76.88 ± 2.36 (-82.07, -71.69) | t(20.66)=0.516 | 0.611 |
| Input resistance (MΩ) | 67.81 ± 9.66 (46.94, 88.68) | 49.82 ± 5.87 (36.90, 62.74) | t(20.99)=1.591 | 0.127 |
| Rheobase (pA) | 341.9 ± 31.4 (274.1, 409.8) | 381.8 ± 33.2 (308.8, 454.7) | t(23.55)=0.872 | 0.392 |
| AP half-width (ms) | 1.369 ± 0.070 (1.218, 1.520) | 1.209 ± 0.072 (1.051, 1.368) | t(23.69)=1.589 | 0.125 |
| AHP amplitude (mV) | -20.77 ± 1.17 (-23.31, -18.24) | -22.59 ± 1.97 (-26.92, -18.26) | t(18.27)=0.792 | 0.439 |
| Accommodation ratio (ISI<sub>late</sub>/ISI<sub>initial</sub>) | 1.310 ± 0.060 (1.181, 1.439) | 1.269 ± 0.070 (1.115, 1.423) | t(22.67)=0.446 | 0.660 |
| Mean ISI (ms) | 24.51 ± 1.34 (21.61, 27.40) | 26.98 ± 1.75 (23.13, 30.83) | t(21.46)=1.123 | 0.274 |

**Table 2.** Mixed effects model for F-I curve (Panel B).

| Source | F(df1, df2) | p |
| --- | --- | --- |
| Injected current | F(12,285)=124.6 | <0.0001 |
| Treatment | F(1,24)=1.877 | 0.183 |
| Current × Treatment | F(12,285)=0.954 | 0.493 |

**Supplemental Figure 3.**
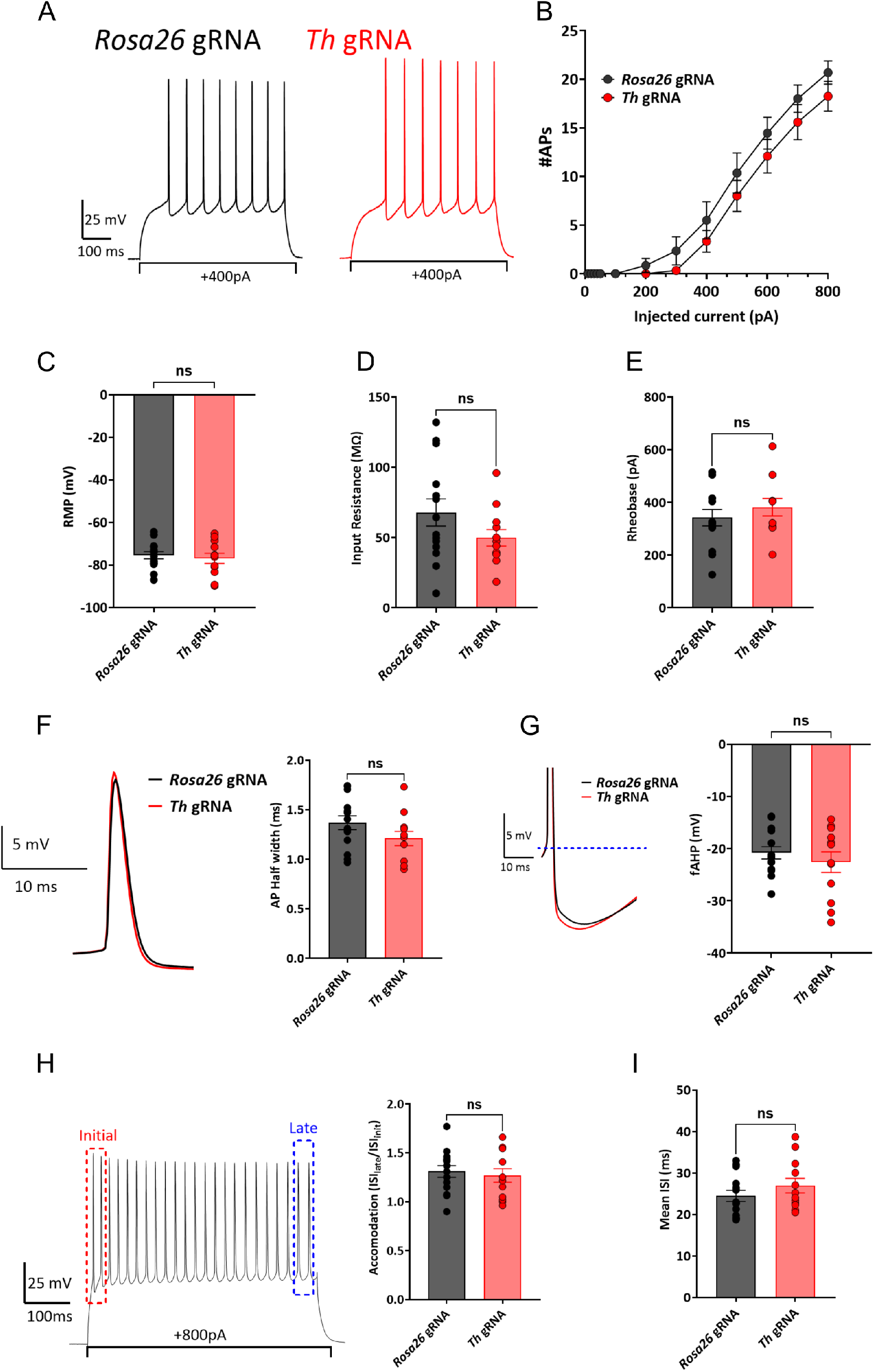
Whole-Cell patch clamp recordings show no significant changes to striatal MSNs after *Th* knockout. Recordings were performed in striatal brain slices from 5 rats. Analyses were performed on 14 neurons from the control hemisphere and 12 neurons from the *Th* gRNA hemisphere. Whole-cell current-clamp recordings revealed no significant differences between *Rosa26* and *Th* KO neurons in number of action potentials (A-B), resting membrane potential (C), input resistance (D), rheobase (E), half-width (F), fast afterhyperpolarization amplitude (G), adaptation ratio (H), or mean interspike interval (I) (all p > 0.12; Table 1). Analysis of firing frequency-current relationships using a REML mixed-effects model demonstrated a significant effect of injected current (F(12,285)=124.6, p<0.0001), but no main effect of treatment (F(1,24)=1.877, p=0.183) and no treatment × current interaction (F(12,285)=0.954, p=0.493), indicating similar excitability across current injection levels in Control and TH KO neurons (Table 2).

